# How do wind speed, release height, seed morphology interact to determine seed dispersal trajectory of *Calligonum* (Polygonaceae) species Wind speed, release height, seed morphology determine seed dispersal trajectory

**DOI:** 10.1101/362434

**Authors:** Quanlai Zhou, Zhimin Liu, Zhiming Xin, Jianqiang Qian, Yongcui Wang, Wei Liang, Xuanping Qin, Yingming Zhao, Xinle Li, Xue Cui, Minghu Liu

**Affiliations:** Institute of Applied Ecology, Chinese Academy of Sciences, 72 Wenhua Road, Shenyang 110016, China; Experimental Center of Desert Forestry, Chinese Academy of Forestry, 1 Tuanjie Road, Dengkou, Bayan Nur 015200, China; University of Chinese Academy of Sciences, 19 Yuquan Road, Beijing 100049, China; Henan Agricultural University, 63 Nongye Road, Zhengzhou 450002, China; Prevention and Quarantine Bureau of Forestry Pest of Liaoning, Shenyang 110036, Liaoning Provence, China

**Keywords:** dispersal distance, dispersal pattern, morphological traits, primary dispersal, trajectory mode, video recording methods, wind tunnel

## Abstract

How seed dispersal trajectory shifts with abiotic and biotic factors and what is the relationship between seed dispersal distance and dispersal trajectory are remain unclear. We used wind tunnel and video camera to track the seed dispersal trajectory of 7 *Calligonum* species with different appendages under the different wind speeds and the release heights. Dispersal trajectories and distances were determined by video analysis and spatial coordinate transformation. Based on perspective principle, 4 modes of trajectories were determined. Wind speed, seed mass and release height were the key factors determining seed dispersal trajectory modes. Release height and wind speed tended to have the strongest explanatory power on seeds with bristles and wings, respectively. Different trajectory modes lead to different dispersal distance, while the same dispersal distance can be the result of different trajectory modes. The proportion of species’ trajectory modes formed its trajectory spectrum. Wind speed tends to have strong influence on light and low-wind-loading seeds, release height tends to have that on heavy and high-wind-loading seeds. Species with high proportion of horizontal projectile and projectile have high dispersal capacity, vice versa. Therefore, trajectory spectrum of a species reveals its primary dispersal strategies and evolutionary consequences.

## 1 Introduction

Exploring the reason why different diaspores have different dispersal distances has been a major challenge for ecologists (Maurer *et al*., 2013; Nathan, 2001; Tackenberg *et al*., 2003; Thomson *et al*., 2011). Seed moving trajectory may explain seed primary dispersal distance (Dauer *et al*., 2006; Jongejans and Telenius, 2001). Determining seed dispersal trajectory and its relevant factors is essential to clarify the mechanisms underlying the structure and dynamics of plant population and community (Nathan, 2001).

Seed dispersal trajectory may be influenced by wind conditions (wind speed and orientation), release height and seed traits such as seed weight, appendage, wind loading and terminal velocity (Savage *et al*., 2014; Thomson *et al*., 2011; Zhu *et al*., 2016). Horizontal wind can disperse seeds far away from their mother plants (Augspurger, 1986; Soons *et al*., 2004; vanDorp *et al*., 1996). Convective updrafts are proposed to promote dispersal distance (Heydel *et al*., 2014; Nathan *et al*., 2002; Soons *et al*., 2004) and change seed dispersal trajectory. So, it is important to quantify the relationship between seed dispersal trajectory and wind conditions.

Release height is considered as an important plant trait affecting seed flying time (Tackenberg, 2003), which is closely related to seed primary dispersal distance and pattern (Thomson *et al*., 2011; vanDorp *et al*., 1996). Seed traits such as seed mass (Augspurger, 1988), terminal velocity (Andersen, 1993) and wind loading (Matlack, 1992) have profound effects on seed dispersal pattern and distance. Seed morphological characters are evolved due to the selection on dispersal ability (Andersen, 1993; Nathan, 2001). Particular seed appendages are generally assumed to help wind-dispersed seeds travel further since they decrease terminal velocity and wind loading but increase the roughness of seed surface (Jongejans and Telenius, 2001; Nathan *et al*., 2011). However, up to date few studies have been conducted on the relationship among release height, seed morphological character, and seed moving trajectory.

In reality, wind conditions are so complicated that it is almost insurmountable to measure (Jones and Muller-Landau, 2008; Jongejans and Telenius, 2001). The wind tunnel, manufacturing controllable wind conditions, is necessary for quantifying the primary seed dispersal distance and distribution pattern (Dauer *et al*., 2006; Zhu *et al*., 2016). It can also be used to determine the dispersal trajectory if special facilities and techniques could be applied.

In this study, wind tunnel and video camera were used to track the seed dispersal trajectory of 7 *Calligonum* (*Polygonaceae*) species with different appendages (wings, bristles, membranous-saccate and wings+thorns) under the wind speed of 4, 6, 8 and 10 m s^−1^ and the release height of 0.2, 0.4, 0.6, 0.8 and 1.0 m. Dispersal trajectories were determined by video analysis and coordinate transformation. Redundancy analyses (RDA) was used to explore the relationship between seed moving trajectory and wind speed, release height and seed morphological traits. The objectives of this study were to answer the following questions: 1) how did seed dispersal trajectory shift with wind speed, release height and seed morphological traits? 2) what is the relationship between seed primary dispersal distance and its dispersal trajectory? We especially tested the hypothesis: 1) in descending order, the key factors determining seed dispersal trajectory modes are wind speed, seed morphological traits and release height; 2) key factors shaping primary dispersal trajectory are different for seeds with various morphological traits: wind speed is the key factor for the seed with wings, while terminal velocity is the key factor for the seed with bristles; 3) the same dispersal distance may be the results of different trajectories; the straight-line-shaped trajectory tend to show a short dispersal distance, but the curve-shaped trajectory tend to show long dispersal distance.

## 2 Materials and Methods

### 2.1 Seed collection

Seeds of 7 *Calligonum* species (Fig. 1) were collected in Ulanbuh Desert (106°09 ′–107°10 ′E, 40°09 ′–40°57 ′N, 1050 m above sea level) in Inner Mongolia and Tazhong (83°40 ′E, 39°00 ′N, 1099 m above sea level) of Taklimakan Desert Xinjiang, Northwestern China. Four hundred intact seeds were collected for each species from 20 individual plants after seed maturity in natural populations in July and August, 2016. All seeds were air dried and stored in laboratory until experiment.

**Fig. 1.**
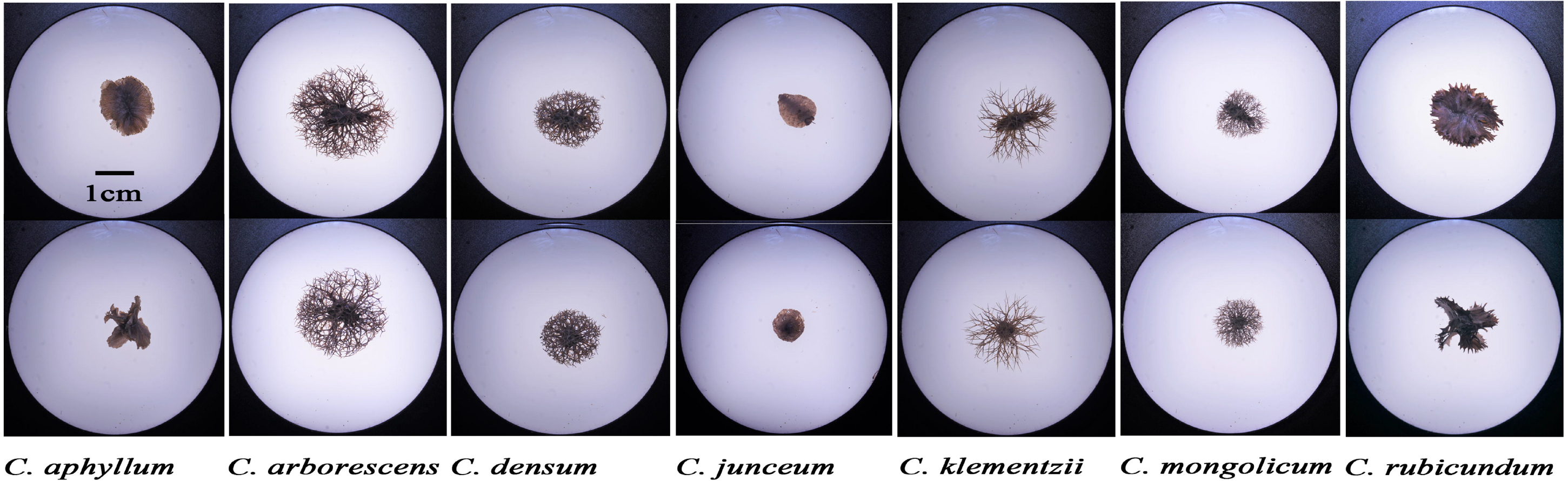
Side and top view of 7 species of Calligonum

### 2.2 Measurement of morphological and aerodynamic traits

Twenty intact seeds were randomly selected to measure the dimension, shape index, mass, projected area, wing loading and terminal velocity. Three dimensions (length, width and height) of each seed were measured using digital caliper (0.01 mm accuracy) (Zhu *et al*., 2016). Seed mass was determined by electronic balance (0.1 mg accuracy). Seeds were scanned and then analyzed by WinSEEDLE (Regent Instruments Inc., Canada) image analysis system to measure projected areas (PA) (Zhu *et al*., 2016). The wing loading (WL) was calculated as the ratio of seed mass to projected area (Greene and Johnson, 1997; Howlett, 1995; Matlack, 1987). The shape index (SI) was calculated following Thompson et al. (1993). The terminal velocity (TV) was determined using an apparatus described by Zotz et al. (2016) (Table 1).

**Table 1.**
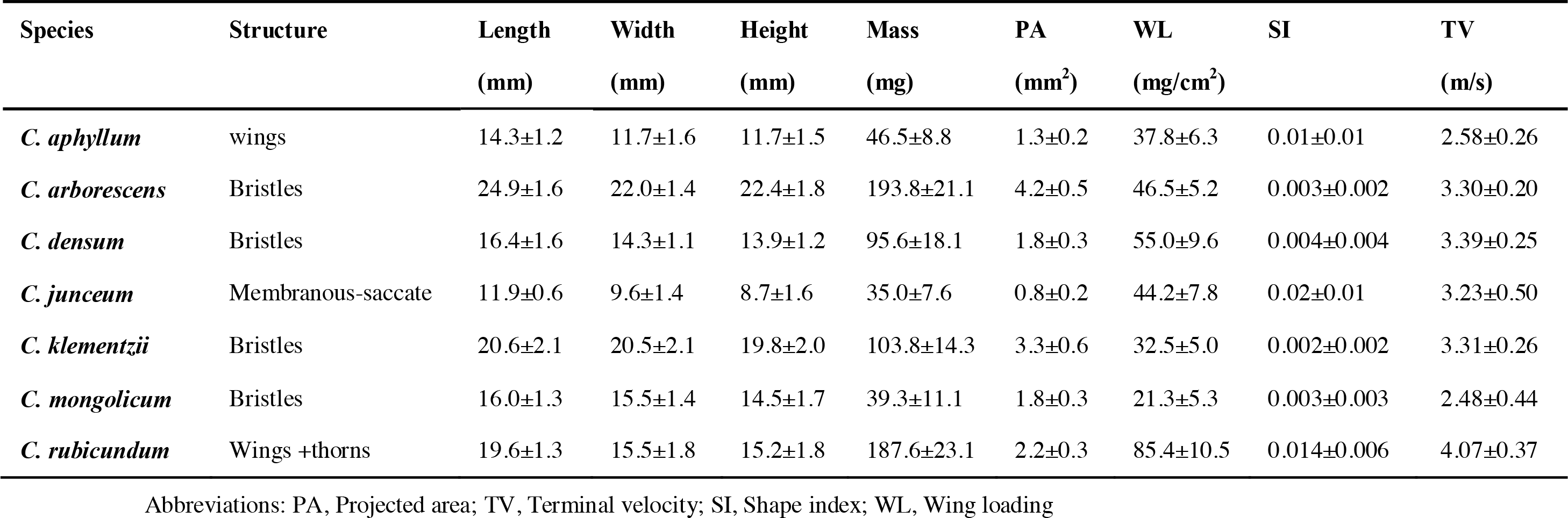
Morphological and aerodynamic characteristics of the seeds in the 7 studied *Calligonum* species (Mean±SD)

### 2.3 Wind-tunnel experiments

A wind tunnel was used to measure the primary dispersal distance and trajectory of *Calligonum* seeds (Zhu *et al*., 2016). A square grids scale (N) was drawn on the wind tunnel side wall (Fig. 2). Markers (M) corresponding to the square grids were drawn on the opposite wall at the bottom of observation window. Marking lines (P) were drawn in order to measure the position of falling seed. A pitot tube (A) suspended from the ceiling and a differential pressure transmitter (B, Dwyer Instruments Inc., Indiana, USA) were used to monitor the wind speed at the test section. Wind speed was measured at the height of 1 m above the ground. Seeds were released via a seed releaser (C) inserted through the middle of the tunnel ceiling. Seeds were put into the top inlet (2 m in length, 6 cm in diameter) of the releaser. A baffle plate (E), controlled by a wrench (D), was covered on the outlet of the releaser in order to keep the seed motionless. The seed release height (H) was the distance from the ground to the outlet.

Twenty treatments in total were designed including four wind speeds (4, 6, 8 and 10 m s^−1^) and five release heights (0.2, 0.4, 0.6, 0.8 and 1.0 m). Twenty seeds (replications) for each species were tested for each treatment. In total, 2800 seeds were released during the wind tunnel experiment. A video camera (G) (GC-P100AC, JVC, Japan) was used to record the dispersal processes using the speed of 50 frames per second (fps) (Fig. 2). The shooting distance was adjusted according to wind speed and release height in order to obtain the overall view of seed dispersal processes.

**Fig. 2.**
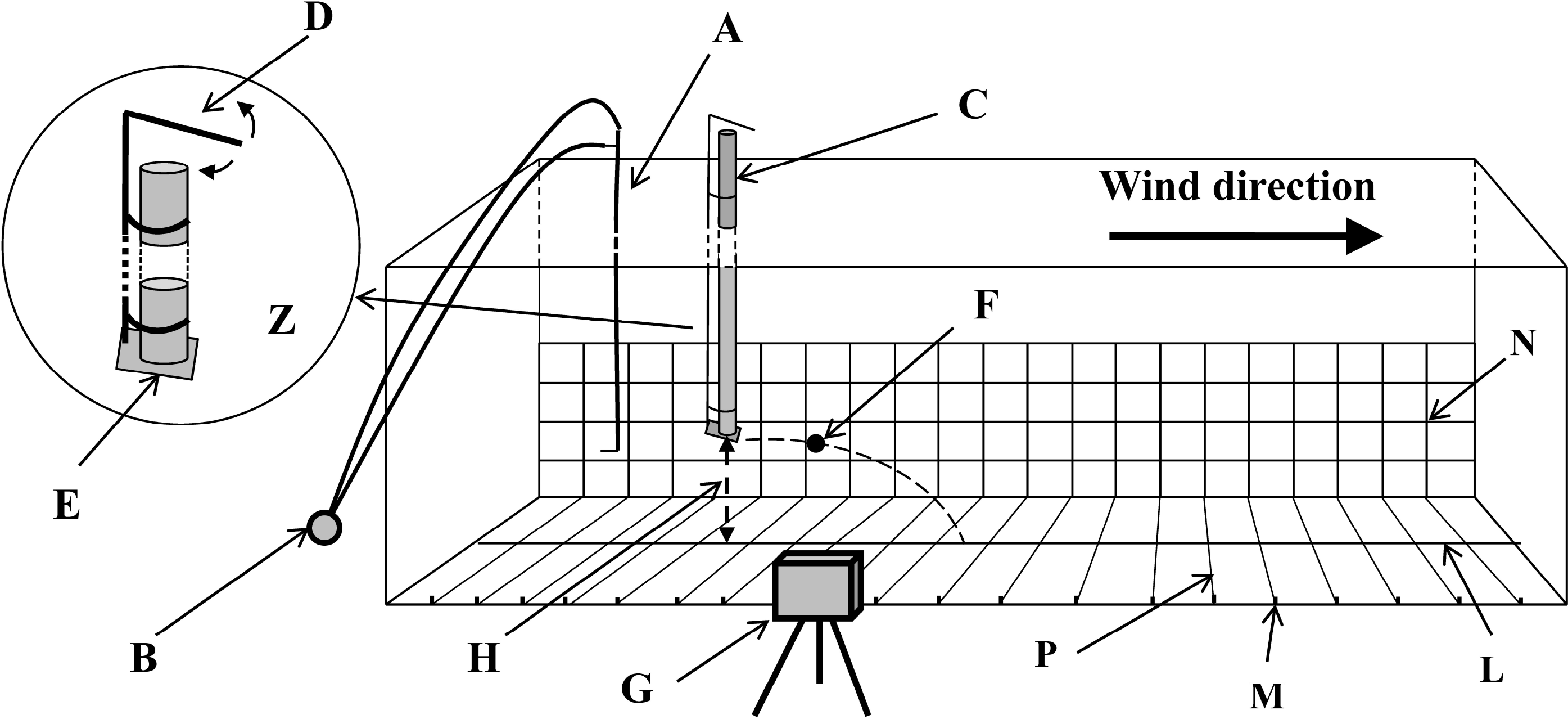
Sketch map of recording seed trajectory and dispersal distance in wind tunnel. A: a pitot tube; B: differential pressure transmitter; C: a seed releaser; Z: detailed structure of the releaser; D: a release controller with a wrench; E: a baffle plate controlled by the wrench; F: a seed released from the releaser; H: release height; G: a camera; M: a marker; L: the centre line of the tunnel ground; N: square grids scale; P: a marking line.

### 2.4 Determination of trajectory and dispersal distance

Videos were analyzed by QuickTime player (Apple Macintosh, version 7.79.80.95, 2016) to determine the seed position and dispersal distance. As the video captured by camera was a 3-dimensional scene, coordinates transformation was conducted to obtain projected coordinates on tunnel wall based on perspective principle (Fig. 3). Every frame of the video was printed on a paper, then two marking lines GH and JK were drawn on the paper intersecting at the point V according to the perspective principle, i.e., the vanishing point. Line HK was an intersection line of side wall and the ground. The point A was a position of falling seed in the picture. Vertical line was drawn from the point A intersecting with the central line LM at point B. Point C was an intersection between line BV and HK. A vertical line was drawn from the point C intersecting with the line AV at point P_i_. The P_i_ (x_i_, y_i_) was the projected coordinates on the scale (Fig. 3). By the same token, successive projected coordinates of every video frame were obtained. A projected trajectory of seed was drawn from the release point P_o_ (x_o_, y_o_) to the fall point P_t_ (x_t_, 0) on the paper. The dispersal distance DP_t_ is (x_t_–x_o_) cm, where the x_t_, and x_o_ is the abscissa of P_t_ and P_o_ (Fig. 3).

**Fig. 3.**
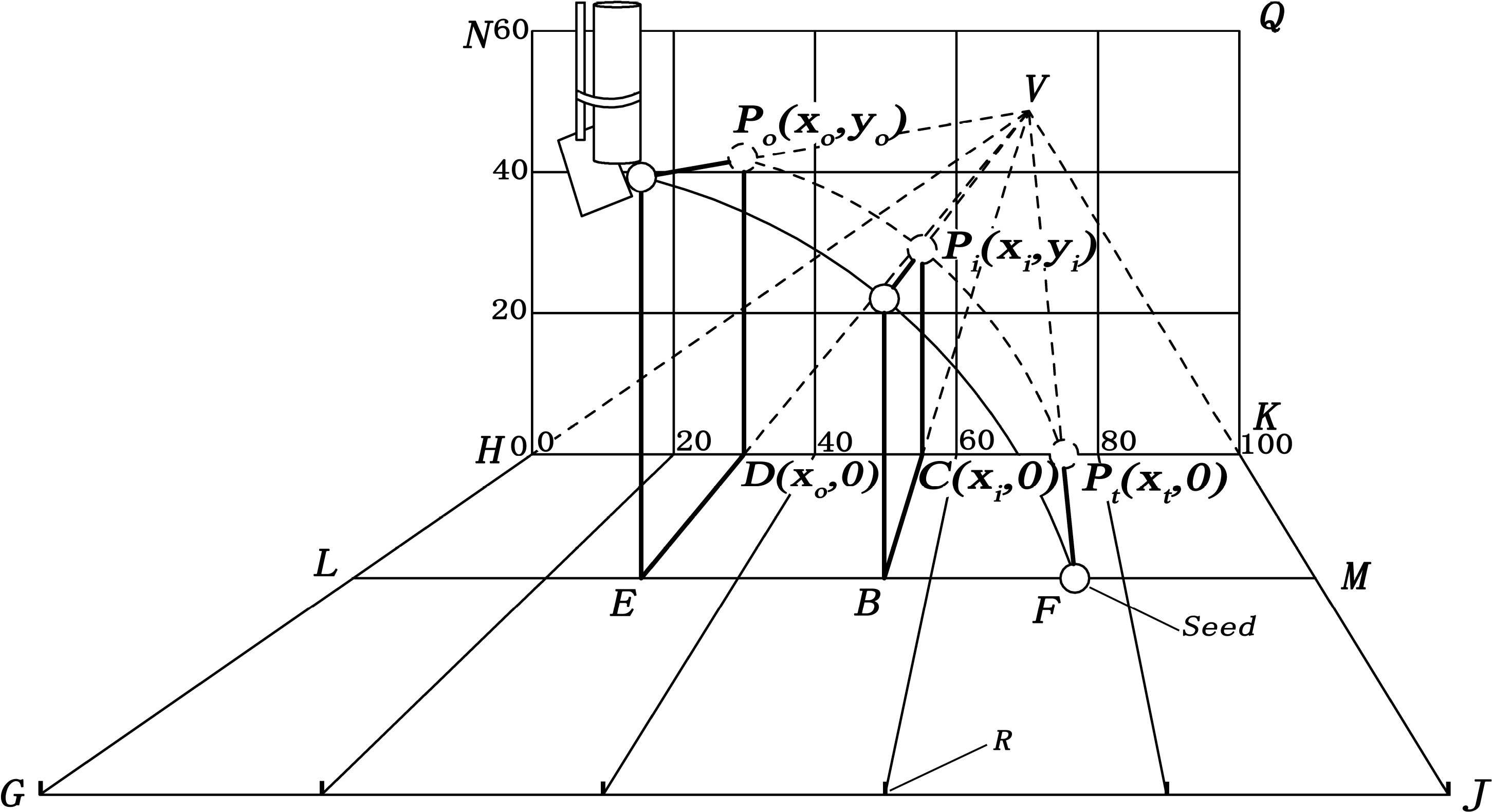
Sketch map of coordinate transformation of seed position from a 3-dimensional scene into a 2-dimensional surface based on perspective principle. NHKQ: square grids scale on the wall of wind tunnel; HGJK: ground of wind tunnel; HK and GH: intersection lines of the side wall and the ground; LM: the centre line of the ground; GH and JK: two marking lines; V: vanishing point, extended lines of GH, JK and their parallel lines intersected at the point; O: seeds release point; A: any point in seed dispersal trajectory; F: seed fall point; E and B: projected point of O and A on the ground; D (x_o_, 0), C (x_i_, 0) and P_t_ (x_t_, 0): intersection points of VE, VB and VF with HK; P_o_ (x_o_, y_o_), P_i_ (x_i_, y_i_) and P_t_ (x_t_, 0): projected points of point O, A and F on the scale; R: a marker.

### 2.3 Data analysis

Redundancy analyses (RDA) was conducted by using Canoco 5.0 (version 5.0, Microcomputer Power, Ithaca, NY, USA) (Tackenberg, 2003) to assess the explanatory power of wind speed (WS), appendage structure (AS), shape index (SI), seed mass (SM), wing loading (WL), terminal velocity (TV) and release height (RH) to the trajectory modes of the 7 *Calligonum* species. Since four types of appendage structures and trajectory modes are not scaled attributes, we defined 4 variables for the occurrence of appendage structures, AS_1_, AS_2_, AS_3_ and AS_4_, with the property that AS_i_=1 if the appendage structure belongs to wings, bristles, membranous-saccate and wings +thorns, otherwise AS_i_=0. For same token, we defined 4 variables for the occurrence of trajectory modes: concave upward, straight line, horizontal projectile and projectile, as TM_1_, TM_2_, TM_3_ and TM_4_. Each explanatory power was tested to assess the contribution of various abotic and botic factors to the variation in dispersal trajectory shifts by using forward-selection procedures.

Hierarchical clustering analysis was used to classify the dispersal distance of 7 species. A dendrogram was drawn by ward’s method after transforming values of dispersal distance into standardized Z scores. The hierarchical clustering analysis, dendrogram plot and descriptive statistics were analysed using PASW Statistics Software version 18.0 (IBM Corp., Armonk, New York, USA). Box plot, curve fit and scatter plot was drawn using SigmaPlot version 10.0 (Systat Software, Inc., USA).

## 3. Result

### 3.1 Trajectory modes of 7 *Calligonum* species under different release heights and wind speeds

Four modes of moving trajectory, i.e., concave upward, straight line, horizontal projectile and projectile, were tracked under different release heights and wind speeds. The concave upward had the largest launching angle between horizontal and moving direction. The straight line had a constant and smaller included angle than that of concave upward. The horizontal projectile had the smallest launching angle (approximate to 0). The projectile had a launching angle above the horizontal direction (Fig. 4).

**Fig. 4.**
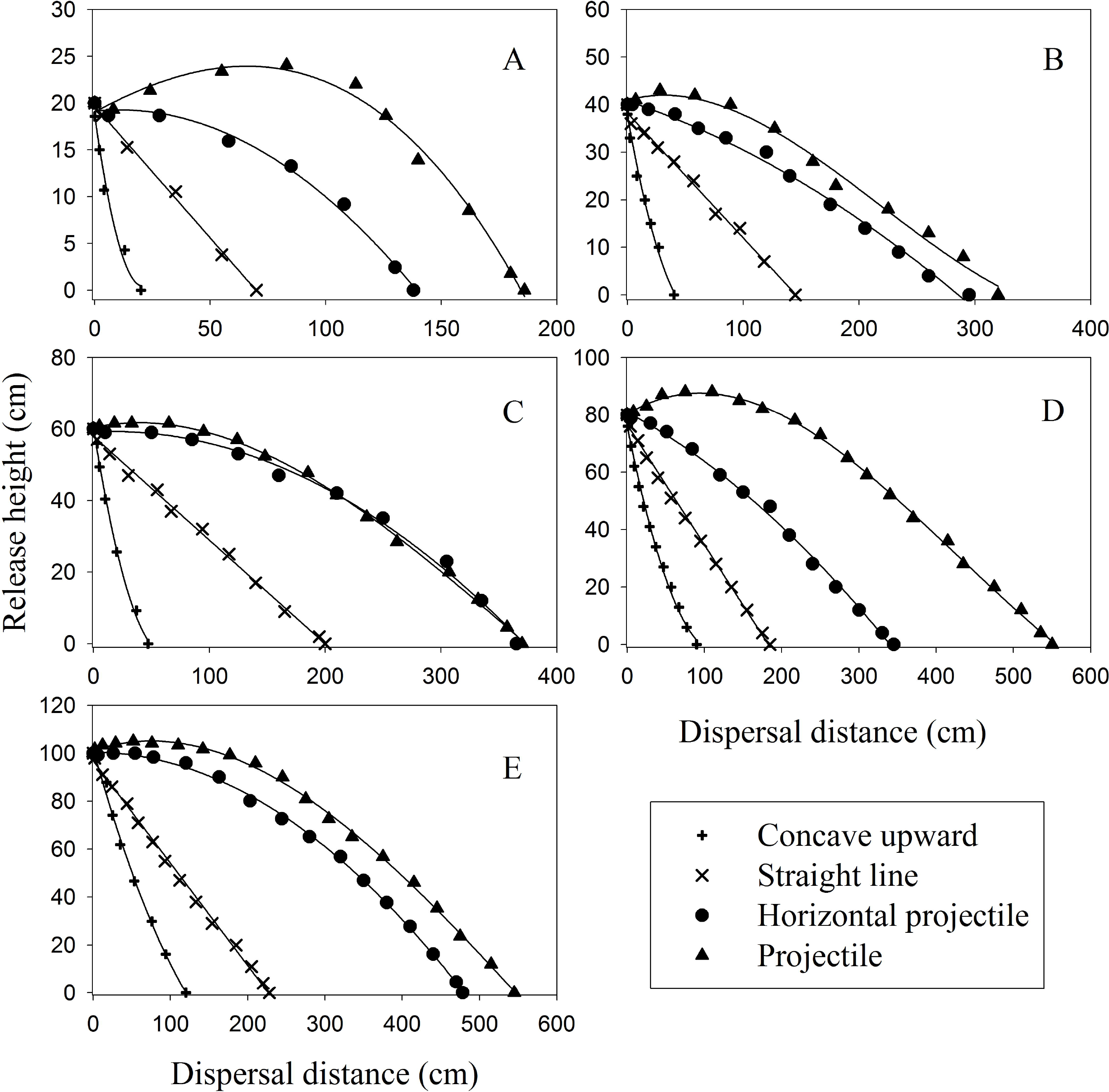
Trajectory modes of primary seed wind dispersal of 7 *Calligonum* species under release height of 20 cm (A), 40 cm (B), 60 cm(C), 80 cm (D) and 100 cm (E) under the wind speed of 4, 6, 8 and 10 m s^−1^.

### 3.2 Relationship between seed dispersal distance and dispersal trajectory mode

The dispersal distance increased from concave upward, straight line, horizontal projectile to projectile. The concave upward trajectory had the dispersal distance ranging from 10 cm to 260 cm (median 45 cm). The dispersal distance of straight line mode ranged from 6 cm to 480 cm (median 105 cm). The horizontal projectile mode dispersed 70 cm to 600 cm in distance (median 260 cm) and the projectile mode from 128 cm to 750 cm (median 337 cm) (Fig. 5).

**Fig. 5.**
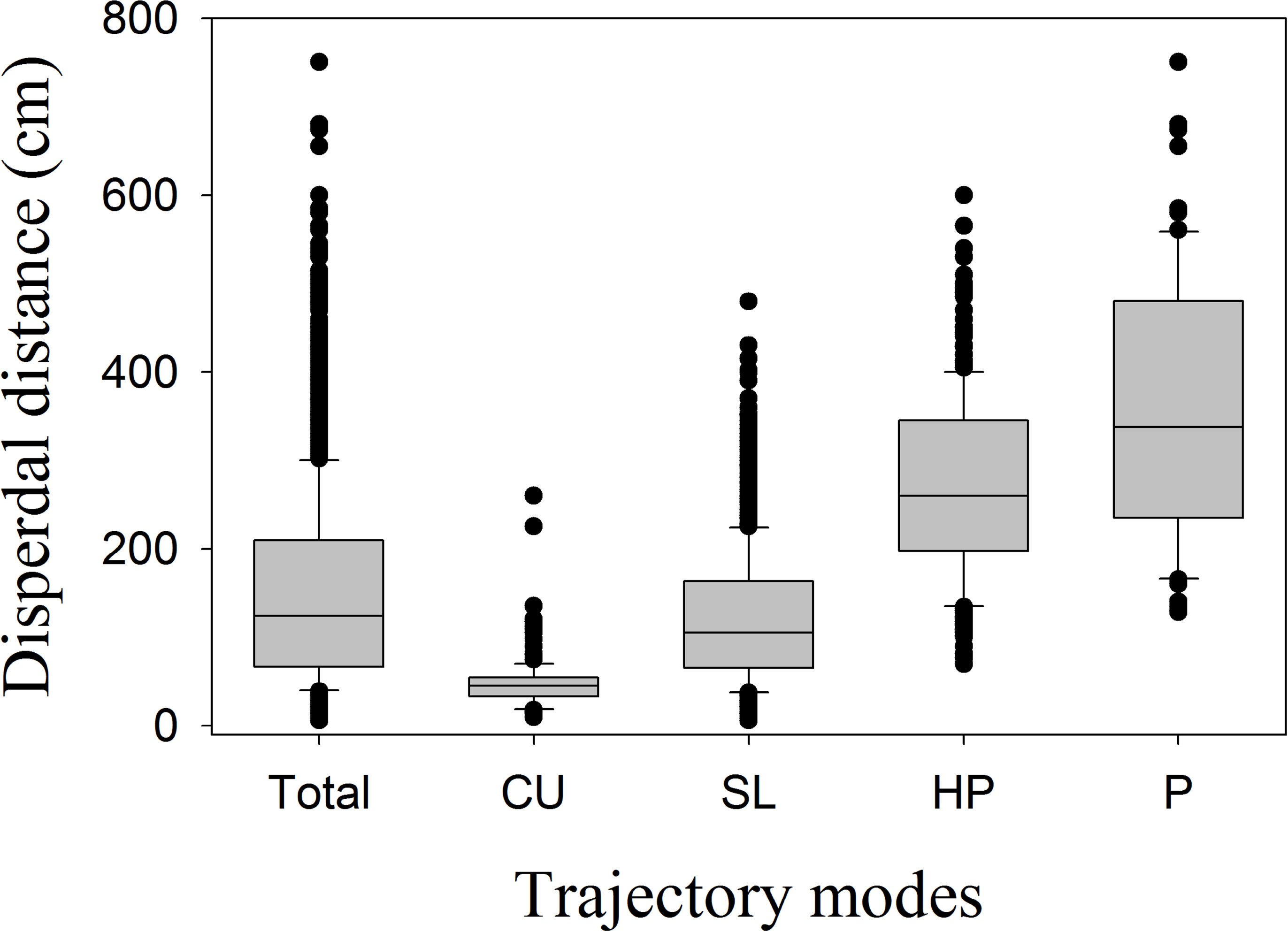
Relationship between trajectory modes and primary dispersal distance. Where, CU: Concave upward; SL: Straight line; HP: Horizontal projectile; P: Projectile.

### 3.3 Occurrence probability of 4 trajectory modes under 4 wind speeds and 5 release heights

Four modes of trajectory shifted with a probability of occurrence under each wind speed and release height. With the increase in wind speed, the concave upward mode showed random occurrence probability, ranging from 17.0%-32.0% (Fig. 6 A). The straight line mode had a decreased probability, ranged from 31.6% to 18.3% (Fig. 6B). The projectile and projectile modes showed increased probability, from 1.1% to 47.1% and from 2.3% to 50.6%, respectively. The accumulative probability of these two modes was more than 80% when the wind speed was above 8 m s^−1^ (Fig. 6 C and D).

**Fig. 6.**
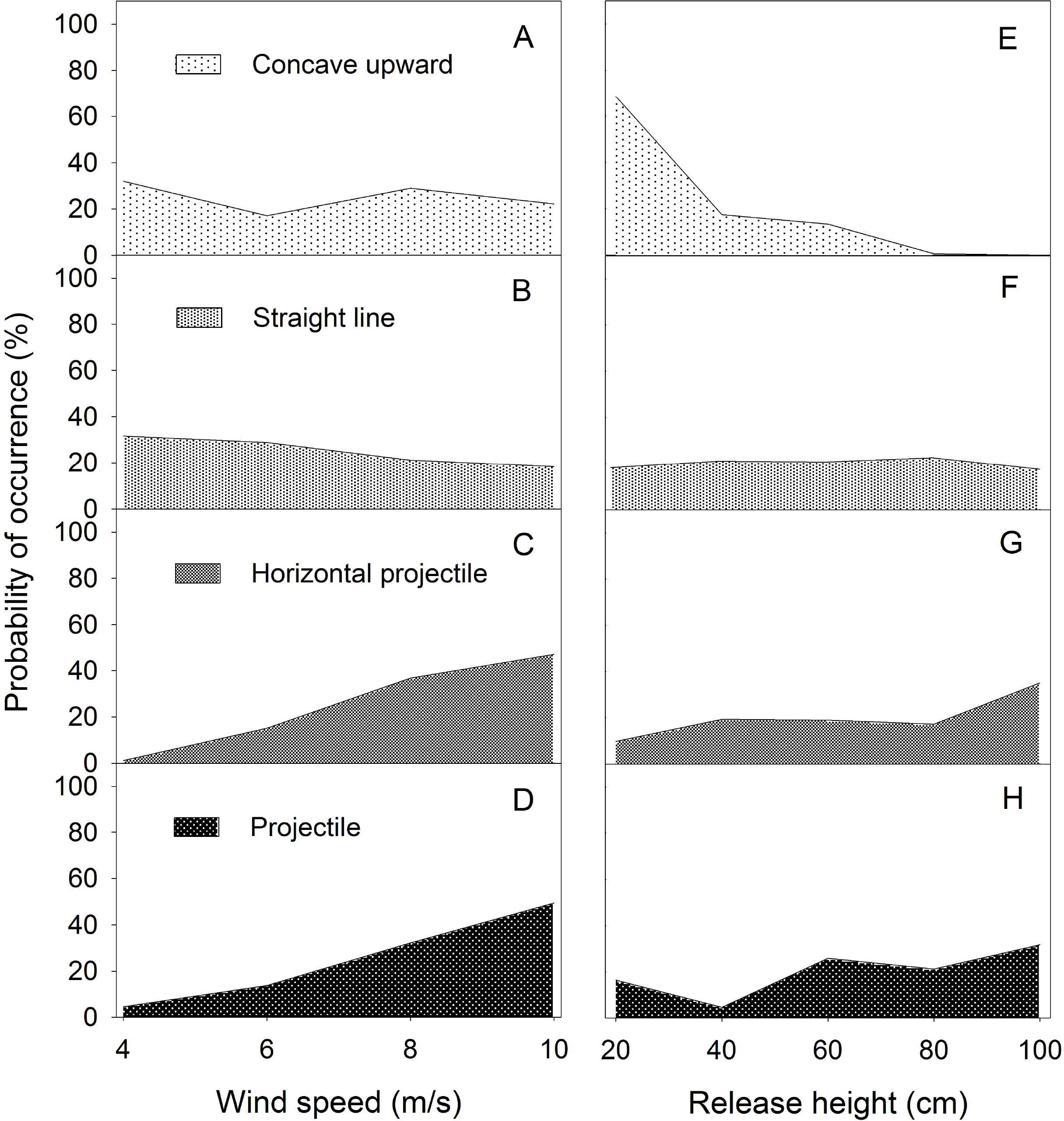
Probability of the occurrence of 4 trajectory modes under 4 wind speeds and 5 release heights. The left panels (A~D) show probability of the occurrence of 4 trajectory modes under 4 wind speeds. The right panels (E~H) show probability of the occurrence of 4 trajectory modes under 5 release heights.

With the increase in release height, the concave upward mode significantly decreased in occurrence probability, from 68.5% to 0, occurred mainly within 20-40 cm (Fig. 6E). The occurrence probability of the straight line mode fluctuated slightly, from 17.4% to 22.6% (Fig. 6 F). The horizontal projectile and projectile modes showed increasing tendency from 9.6%-34.9% and 4.7%-31.7%, respectively. Each of these two modes had the occurrence probability higher than 75% within the release height 60 cm to 100 cm (Fig. 6 G and H).

### 3.4 Clustering of dispersal distance of 7 *Calligonum* species

The 7 species could be classified into 3 groups according to the dispersal distance (Fig. 7). Group I included *C. arborescens*, *C. densum*, and *C. klementzii*, with bristles, big mass and high terminal velocity (Table 1). Mean of dispersal distance was 124.3 cm (standard deviation, 84.3) ranging from 10 cm to 395 cm (median 104.5 cm). Group II included *C. mongolicum* and *C. rubicundum* with bristles and wings+thorns, respectively. *C. mongolicum* had small mass and terminal velocity, while *C. rubicundum* had big mass and terminal velocity (Table 1). Mean of dispersal distance was 151.7 cm (standard deviation 108.8) ranging from 6 cm to 650 cm (median 124.0 cm). Group III included *C. aphyllum* and *C. junceum* with small mass and terminal velocity. *C. aphyllum* had wings and *C. junceum* had balloon-like membranous-saccate. Mean of dispersal distance was 181.0 cm (standard deviation 120.8) ranging from 18 cm to 750 cm (median 150.0 cm).

**Fig. 7.**
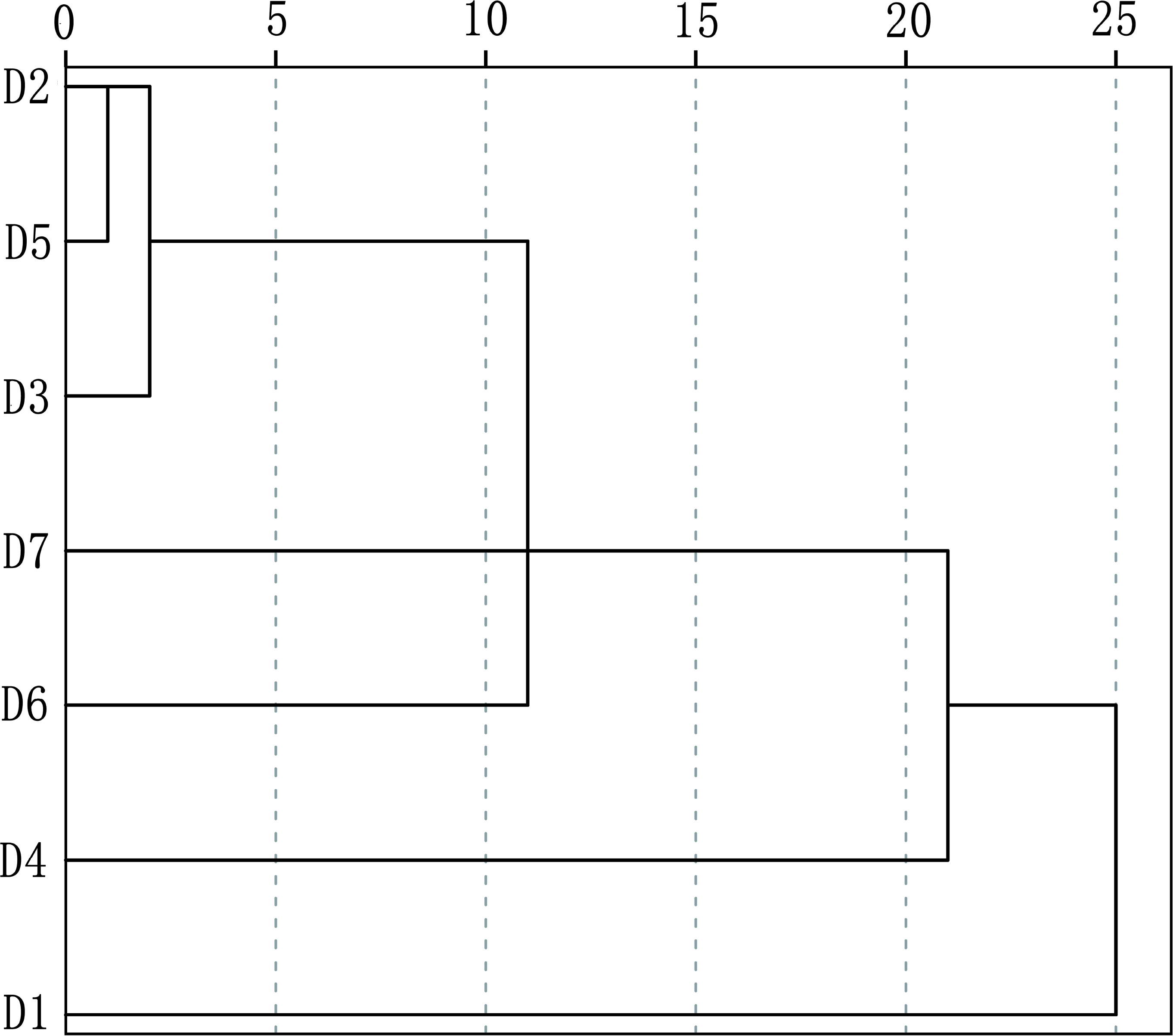
The clustering of primary dispersal distance of 7 *Calligonum* species using hierarchical method (D1) *C. aphyllum;* (D2) *C. arborescens;* (D3) *C. densum;* (D4) *C. junceum;* (D5) *C. klementzii;* (D6) *C. mongolicum;* (D7) *C. rubicundum*

### 3.5 Distribution of trajectory modes in 7 *Calligonum* species

In group I, the modes of concave upward trajectory (>10%) and straight line trajectory (>75%) accounted for more than 90% of total trajectories, while the modes of horizontal projectile trajectory (<7.5%) and projectile trajectory (not found in the three species) was less than 10% (Table 2). In group II, the concave upward and straight line modes in two species occupied 58-85% of total trajectories and the horizontal projectile and projectile 13.8%-41% (Table 2). In group III, the concave upward and straight line modes in two species were less than 65% of the total trajectories. The horizontal projectile and projectile modes made up more than 35% of the total trajectories (Table 2).

**Table 2.** Number and proportion of 4 trajectory modes in each species.

| Group | Species | Concave<br>upward | Straight<br>line | H-projectile | Projectile |
| --- | --- | --- | --- | --- | --- |
| Group I | *C. arborescens* | 50 (12.5%) | 345 (86.3%) | 5 (1.2%) | 0 |
|  | *C. densum* | 82 (20.5%) | 300 (75.0%) | 18 (4.5%) | 0 |
|  | *C. klementzii* | 41 (10.2%) | 329 (82.3%) | 30 (7.5%) | 0 |
| Group II | *C. mongolicum* | 5 (1.2%) | 227 (56.8%) | 162 (40.5%) | 6 (1.5%) |
|  | *C. rubicundum* | 10 (2.5%) | 335 (83.7%) | 55 (13.8%) | 0 |
| Group III | *C. aphyllum* | 5 (1.2%) | 204 (51.0%) | 149 (37.3%) | 42 (10.5%) |
|  | *C. junceum* | 1 (0.3%) | 252 (63.0%) | 111 (27.7%) | 36 (9.0%) |

We defined the proportion of trajectory modes in a species as the trajectory spectrum. In group I, there is higher percentage of concave upward and straight line, and low percentage of horizontal projectile and projectile in their spectrum, such as *C. arborescens* and *C. densum* (Table 2). In group III, high percentage of horizontal projectile and projectile occurs in their spectrum, such as *C. aphyllum* (Table 2).

### 3.6 Relationship between wind speed, release height, seed morphological traits, and trajectory modes

The terminal velocity (TV), wings (AS_1_), membranous-saccate (AS_3_), release height (RH) and shape index (SI) were positively linked with the modes of horizontal projectile (TM_3_) and projectile (TM_4_), while negatively linked with the concave upward (TM_1_) and straight line (TM_2_) (Fig. 8 A). The seed mass (SM), TV, bristles (AS_2_), wings+thorns (AS_4_) and wind loading (WL) were negatively linked with horizontal projectile (TM_3_) and TM_4_, and positively linked with TM_1_ and TM_2_ (Fig. 8 A).

**Fig. 8.**
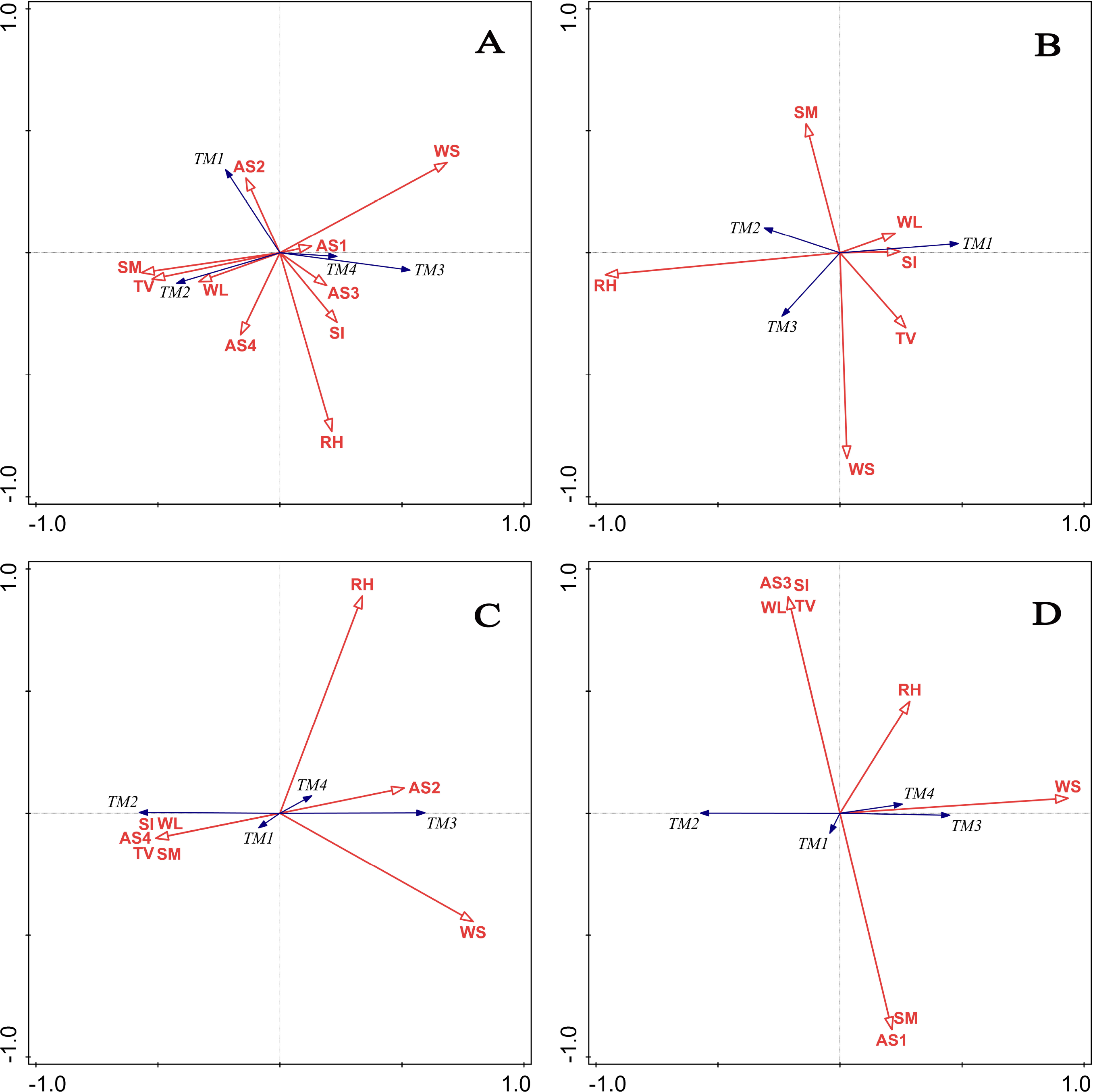
Relationship between wind speed, release height, seed morphological traits, and shifts of trajectory mode of (A) 7 species (in total); (B) 3 species in group one; (C) 2 species in group two; (D) 2 species in group three. TM: trajectory mode; TM_1_: concave upward, TM_2_: straight line; TM_3_: horizontal projectile; TM_4_: projectile; SM: seed mass (mg); TV: terminal velocity (m/s); WL: wing loading (mg/cm^2^); SI: shape index; RH: release height (cm); WS: wind speed (m/s); AS: appendage structure; AS_1_: wings; AS_2_: bristles; AS_3_: membranous-saccate; AS_4_: wings+thorns. Explanatory factors are indicated by red lines with hollow arrows. Trajectory modes are showed by blue lines with solid arrows.

For all 7 species, the explanatory power on the trajectory modes (TM) in descending order was WS, SM, RH, TV, AS_4_, SI and WL (Table 3). The WS has the most explanatory power, explained 9.2% (*p*=0.002) of variation in trajectory modes. The SM has the second explanatory power, explained 6.1% of the variation. The other factors in descending order were RH, TV, AS and SI as well as WL, totally explained 6.3% of the variation (Table 3).

**Table 3.** Explanations and relative contributions of wind speed, release height and seed morphological traits to trajectory modes of 7 *Calligonum* species

| Species | Explanatory factors | Explanation (%) | Contribution (%) | F value | *p* value |
| --- | --- | --- | --- | --- | --- |
| All 7 species | WS | 9.2 | 42.4 | 283.0 | 0.002 |
|  | SM | 6.1 | 28.1 | 201.0 | 0.002 |
|  | RH | 2.2 | 10.2 | 75.0 | 0.002 |
|  | TV | 1.9 | 8.9 | 36.4 | 0.002 |
|  | AS<sub>4</sub> | 1.1 | 4.9 | 68.0 | 0.002 |
|  | SI | 0.7 | 3.4 | 26.4 | 0.002 |
|  | WL | 0.4 | 1.7 | 12.8 | 0.002 |
|  | Total (%) | 21.6 | 99.6 |  |  |
| Group I | RH | 13.4 | 84.1 | 186.0 | 0.002 |
|  | TV | 1.2 | 7.5 | 16.7 | 0.002 |
|  | WS | 1.0 | 6.6 | 14.8 | 0.002 |
|  | SM | 0.3 | 1.9 | 4.3 | 0.028 |
|  | Total (%) | 16.0 | 100 |  |  |
| Group II | WS | 20.3 | 62.7 | 203.0 | 0.002 |
|  | SI | 8.4 | 25.9 | 93.5 | 0.002 |
|  | RH | 3.7 | 11.4 | 43.5 | 0.002 |
|  | Total (%) | 32.4 | 100 |  |  |
| Group III | WS | 20.3 | 87.2 | 204.0 | 0.002 |
|  | RH | 1.9 | 8.2 | 19.6 | 0.002 |
|  | SI | 1.1 | 4.6 | 11.2 | 0.002 |
|  | Total (%) | 23.3 | 100 |  |  |

In group I, RH, TV, WS and SM were the significant explanatory variables (*p*=0.002), explained 16.0% of the variation in trajectory modes (Table 3). The RH was positively linked with TM_2_, TM_3_ while negatively linked with TM_1_. The TV was positively linked with TM_1_ and negatively linked with TM_2_. The WS was positively linked with TM_3_ and negatively linked with TM_2_. The SM was positively linked with TM_2_ and negatively linked with TM_3_ (Fig. 8 B). The RH was the most significant factor explained 13.4% (*p*=0.002) of the variation in trajectory modes. While TV, WS and SM only explained 2.5% (Table 3).

In group II, WS, SI and RH were the significant explanatory variables (*p*=0.002), explained 32.4% of the variation in trajectory mode shifts (Table 3). The WS and RH were positively linked with TM_3_ and TM_4_ and negatively linked with TM_1_ and TM_2_. The SI was positively linked with TM_1_ and TM_2_ and negatively linked with TM_3_ and TM_4_ (Fig. 8 C). The WS was the key factor, explained 20.5% of the variation. The SI and RH explained 12.1% (Table 3).

In group III, the significant explanatory variables (*p*=0.002) were WS, RH and SI, explained 23.3% of the TM variation (Table 3). The WS and RH were positively linked with TM_3_ and TM_4_ and negatively linked with TM_1_ and TM_2_. The SI was positively linked with TM_2_ and negatively linked with TM_1_, TM_3_ and TM_4_ (Fig. 8 D). The WS was the key factor, explained 20.3% of the variation in trajectory modes. The RH and SI explained 3.0% in total (Table 3).

## 4. Discussion

### 4.1 Contributions of wind speed, release height and morphological traits to trajectory modes

Our results showed that wind speed had the strongest effect on trajectory modes. The finding here support the general perception that wind is the key factor determining seed dispersal trajectory. The trajectory modes might be affected by not only horizontal wind but also turbulent fluctuations in vertical updrafts. Previous studies demonstrated that horizontal wind is an important environmental factor prompting the lateral dispersal (Horn *et al*., 2001; Travis *et al*., 2010). Vertical updrafts are generated by a shearing action of the horizontal wind (Greene and Johnson, 1995; Levine and Murrell, 2003; Soons *et al*., 2004). Fluctuations of shear-induced updrafts would increase with increasing horizontal wind speed (Greene, 2005). Thus, the trajectory modes might be strongly influenced by the fluctuations of updrafts with wind speed. Surprisingly, seed mass has the second explanatory power on trajectory modes though the seeds were selected within the same genus, *Calligonum*, which have little variation in seed mass among species. The high explanatory power of seed mass could be due to the fact that seed mass could influence lateral movement and rate of descent (Matlack, 1987, 1992). The findings are in consistent with the perception that small seeds disperse better than large seed because larger seed mass is poorly accelerated and dispersed by wind (Greene and Johnson, 1993; Thomson *et al*., 2011). Different from our expectations, release height has the third explanatory power on trajectory modes instead of having the least effect on it. The reason might be that release height is strongly correlated with falling time of seed (Savage *et al*., 2014). The falling time can change horizontal and vertical speed of seeds, and thus affect the trajectory modes.

The other seed morphological traits such as terminal velocity, appendage structure, shape index and wing loading explained 4.1% (accounting for 19.0% of total explanation) of the variation in trajectory modes. One possible reason is that these factors have low variation across species, consequently contributing low variation to the trajectory modes.

In this study, the redundancy analysis showed that the total explanation power of abiotic and biotic factors on the variation in trajectory modes is from 16.0% to 32.4%. There are two possible reasons for the low explanatory power occurred in the experiment. Firstly, the occurrence of trajectory modes, such as straight line, varies little with wind speed and release height, which decreases explanatory power of independent variable on shifts of dispersal trajectory. Secondly, the wind loading may be not appropriate for spherical seeds with complicated structures because its projected area might be miscalculated.

### 4.2 Effects of seed appendages on dispersal trajectory modes and spectrum

Appendages of wind-dispersed seeds are designed for slowing descent rate by lowering wing loading or increasing the roughness of seed surface and the chance of exposure to wind (Andersen, 1993; Augspurger, 1986; Nathan *et al*., 2011). Our findings suggest that the key factors shaping primary dispersal trajectory might be different for seeds with different morphological traits. The redundancy analysis showed that the key factor is not terminal velocity but release height for seeds with bristles. However, for seeds with wings, the key factor is wind speed. There are two possible reasons. Firstly, seeds with bristles have high wind loading because the bristles have so low projected area that low drag force is exerted by wind speed. Thus, the trajectory modes are dependent on the release height rather than the terminal velocity or wind speed. Secondly, wings decrease wing loading and obtain more drag force from wind speed. So, wind speed has much effect on the trajectory modes of seeds with wings.

Seed appendages have profound influence on the trajectory spectrum of species. The trajectory spectrum is different between seeds with bristles and seeds with wings. The proportion of concave upward and straight line for seeds with bristles (e.g. *C. arborescens*) are higher than that for seeds with wings (e.g. *C. aphyllum*). Conversely, the proportion of horizontal projectile and projectile for seeds with bristles are lower than that for seeds with wings.

### 4.3 The relationship between seed dispersal distance and trajectory modes

Previous studies demonstrated that wind speed is the most important determinant of seed dispersal distance (Soons *et al*., 2004; Zhu *et al*., 2016). The findings showed that the trajectory of concave upward and straight line tend to have short dispersal distance because wind speed has little influence on the occurrence of these two trajectories. And, concave-upward tend to have the shortest distance because it mostly occurs under the release heights below 40 cm which shortens the dispersal distance. However, the trajectory of horizontal projectile and projectile tend to have long dispersal distance because their occurrence probabilities are greatly affected by wind speed, and mostly occurred under wind speed over 8 m s^−1^.

Obviously, different trajectory modes have different ranges of dispersal distance. However, the studies have adequately support the hypothesis that the same dispersal distance may be the results of different trajectory modes because the ranges of dispersal distance overlap each other among different trajectories, for example the overlap of the straight line (ranged from 6 cm to 480 cm) and horizontal projectile (70 cm to 600 cm) is from 70 cm to 480cm.

### 4.4 The relationship between seed dispersal capacity and trajectory spectrum

Previously studies showed that seed dispersal distance embody dispersal capacity (Thomson *et al*., 2011). Therefore, the trajectory spectrum of a species can reflect its dispersal capacity. For example, *C. arborescens*, *C. densum* and *C. klementzii* have low dispersal capacity due to the high percentage of concave upward (>10%), low percentage of horizontal projectile (<7.5%) and projectile (0) (Table 2). On the contrary, *C. aphyllum* and *C. junceum* have high dispersal capacity since there are low proportion (<2%) of concave upward and high proportion of horizontal projectile (>25%) and projectile (>9%) in their spectrum (Table 2). The results suggest that the trajectory spectrum of each species might reflect the dispersal strategy due to consequence of natural selection.

## 5. Conclusion

In virtue of the wind tunnel and video recording technique, concave upward, straight line, horizontal projectile and projectile are tracked in the 7 species. Wind speed, seed mass and release height are key factors determining seed dispersal trajectory modes. Release height has main influence on primary dispersal trajectory of seeds with bristles, while wind speed has it on that of seeds with wings. Concave upward and straight line modes tend to exert short dispersal distances, the horizontal projectile and projectile modes tend to exert long dispersal distances. Different trajectory modes lead to different dispersal distances, but the same dispersal distance can be resulted from different trajectory modes. Trajectory spectrum of species not only reveals its dispersal capacity but also reflects its primary dispersal strategies and evolutionary consequences. This study provides critical information for clarifying the mechanisms of seed primary dispersal and predicting seed dispersal pattern.

## Acknowledgements

We thank Zhigang Wang for technical support in wind tunnel experiment and valuable comments on the manuscript, to Junliang Gao, Fengmei Luo, Batu Gegen, Jingbo Zhang, Lu Hai, Yaru Huang, Cheng Ge, Na Duan and Ruibing Duan for assisting with seeds collection and technical support in measurements in Experimental Center of Desert Forestry, Chinese Academy of Forestry. This study was supported by National Natural Science Foundation of China (41571270).

## Authors’ contributions

ZL, ML and QZ conceived the ideas and designed methodology; ZX, WL, XQ and XC collected the data; JQ, YW, YZ and XL analyzed the data; QZ led the writing of the manuscript. All authors contributed critically to the drafts and gave final approval for publication.

